# Stereochemically-aware bioactivity descriptors for uncharacterized chemical compounds

**DOI:** 10.1101/2024.03.15.584974

**Authors:** Arnau Comajuncosa-Creus, Aksel Lenes, Miguel Sánchez-Palomino, Patrick Aloy

## Abstract

We recently presented a set of deep neural networks to generate bioactivity descriptors associated to small molecules (i.e. *Signaturizers*), capturing their effects at increasing levels of biological complexity (i.e. from protein targets to clinical outcomes)^1^. However, such models were trained on 2D representations of molecules and are thus unable to capture key differences in the activity of stereoisomers. Now, we systematically assess the relationship between stereoisomerism and bioactivity on over 1M compounds, finding that a very significant fraction (∼40%) of spatial isomer pairs show, to some extent, distinct bioactivities. We then used these data to train a second generation of Signaturizers, which are now stereochemically-aware, and provide an even more faithful description of complex small molecule bioactivity properties.

## Main text

Small molecules are a great tool to probe biology and, still, the main asset of pharmaceutical companies. The last years have seen a surge of ever more complex biological high-throughput assays involving the use of chemical compounds, and databases committed to gathering bioactivity data associated to small molecules are expanding^2, 3^. Moreover, the widespread availability of computational resources^4^ and artificial intelligence techniques has been pivotal to leverage such amounts of data^5^.

From the computational perspective, small molecules are typically characterized by numerical descriptors encoding physicochemical or topological features^6^. Compounds can be further described using their biological activities (e.g. the targets they interact with), which represents a complementary strategy that extends the small molecule similarity principle beyond conventional chemical properties^7^. Unfortunately, experimental bioactivity data are sparse and only available for a limited set of well-characterized compounds. To overcome these coverage issues, we recently trained a set of Signaturizers that are able to infer bioactivity signatures for any compound of interest, even for those that still remain poorly characterized^1^. However, such predictors are built on 2D representations of molecules and are thus not able to capture subtle, but often meaningful, bioactivity differences between stereoisomers. Indeed, stereochemistry and chirality play pivotal roles in pharmacology^8, 9^, often driving supramolecular recognition processes crucial in drug design. Biological matter is intrinsically chiral (e.g. amino acids)^10^ and stereoisomeric small molecule drugs may exhibit different therapeutic and toxicological effects^11, 12^. For example, the antidepressant Citalopram is administered as a mixture of two enantiomers (i.e. racemate), although only one of them is active^13, 14^. Moreover, the non-active enantiomer is associated with toxic side effects. This is the case of the antiarthritic drug Penicillamine, administered as an enantiomerically pure compound ((S)-Penicillamine) since (R)-Penicillamine acts as a pyridoxine (vitamin B_6_) antagonist and is thus toxic ^12, 15^. We now present novel deep learning models to generate stereochemically-aware bioactivity signatures for any compound of interest, which we call *Signaturizers3D*, and overcome the inherent limitations of our original Signaturizers.

### Systematic quantification of the relationship between stereochemistry and small molecule bioactivity

The first steps in the development of Signaturizers3D were (i) to select a comprehensive database containing detailed bioactivity data for a wide range of chemical compounds, and (ii) within this database, systematically identify groups of stereoisomers to compare their bioactivity profiles and evaluate the ability of Signaturizers3D to distinguish them.

To gather bioactivity data, we used the Chemical Checker (CC), which represents the largest collection of small molecule bioactivity signatures available to date, with experimental information for over 1M compounds^7^. The CC divides data into five levels of increasing complexity, ranging from the chemical properties of compounds to their clinical outcomes. Compound bioactivities are expressed in a vector-like format (i.e. signatures), and the data processing pipeline also includes several steps of increasing level of integration and abstraction: from raw experimental data representing explicit knowledge (type 0 signatures) to inferred representations that leverage all the experimentally determined bioactivities available for each molecule (type III signatures). Thus, we processed the whole CC (i.e. 25 different bioactivity types for ∼1M molecules) to systematically identify groups of stereoisomers that might exhibit distinct bioactivities. In brief, we first identified stereoisomers using their InChIKey strings and we then applied several filters to ensure that the actual differences between compounds were exclusively due to stereochemical variations (see *Supplementary Information* for further details). Then, we selectively removed molecules that were not exhaustively characterized, in order to work with enantiomerically pure compounds and prevent the analysis of results derived from racemic mixtures (**Fig 1a**). We eventually identified 23,830 groups of stereoisomers, involving 57,989 compounds, across the different CC bioactivity spaces. We found most stereoisomeric groups with experimental information in the target binding space (B4) and in the network spaces derived from B4 (i.e. C3-5, **Fig S1**). We thus focused our study on the B4 space, which contains over 600,000 molecules, and we identified 15,370 groups of stereoisomers, involving 32,705 compounds (**Fig 1b**). We then analyzed the binding profiles for all these compounds, and found 6,022 groups that had at least 2 stereoisomers with non-identical binding profiles. We also observed that the majority of the groups (14,181, ∼92%) contained only 2 stereoisomers (**Fig 1c**, top), in 38% of which both compounds showed distinct binding profiles (**Fig 1c**, bottom). Analogously, we identified 562 groups containing 3 stereoisomers: 230 (41%), 195 (35%) and 137 (24%) of them showing 1, 2 and 3 distinct binding profiles, respectively. Finally, we observed that the distribution of Jaccard distances between binding profiles of stereoisomeric groups was skewed towards low values (i.e. more similar profiles) compared with random pairs, while pairs of compounds sharing at least one target were somewhere in the middle (**Fig 1d**). **Fig 1e** shows, as an illustrative example, a group of 3 stereoisomers with non-identical binding profiles, where compounds A and C weakly and strongly bind with the Beta-1 adrenergic receptor (ADRB1; 2^nd^ position in the profile), respectively, while compound B does not bind it. Note that inactive compound-target interactions might be false negatives due to, for instance, a limited sensitivity of the detection methods or non-tested enantiomers.

**Figure 1:**
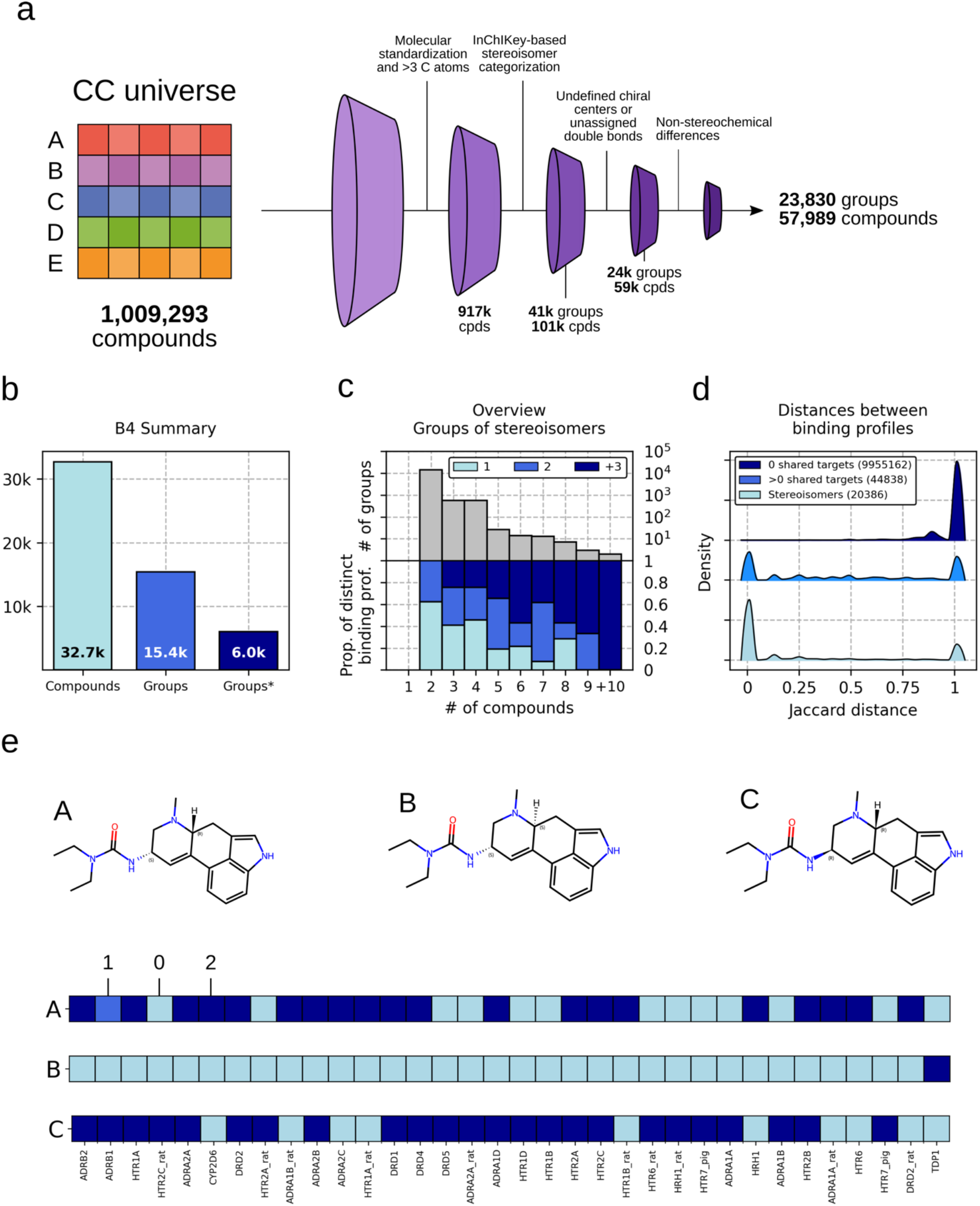
*Stereoisomerism and bioactivity*. **a)** Computational pipeline to identify groups of stereoisomers in the CC chemical universe. **b)** Number of unique stereoisomeric compounds with experimentally identified protein targets in the CC B4 space, number of stereoisomer groups, and number of groups with at least 2 compounds with non-identical binding profiles. **c)** Number of groups (y-axis, top) having the specified number of stereoisomers (x-axis). Proportion of these groups (y-axis, bottom) having the specified number of distinct binding profiles (i.e. ∼60% of the groups of 2 isomers have a unique binding profile). **d)** Distributions of Jaccard distances (binding profiles) between pairs of compounds sharing 0, ≥1 targets and stereoisomer pairs. All distributions are significantly different from each other (Mann-Whitney p-value∼0). **e)** Illustrative example of a stereoisomer group including 3 small molecules with their corresponding target binding profiles, using the annotation of type 0 signatures (i.e. 0: no binding; 1: weak binding and 2: strong binding).

Overall, we observed that most stereoisomer pairs (60.4%) had identical target profiles but, perhaps more interestingly, the remaining 8,081 pairs (39,6%) showed distinct binding against protein targets (**Fig 2a**). Our analyses also showed that CC type III signatures captured differences between stereoisomer pairs (**Fig 2b**). However, these differences were completely missed by the Signaturizers (**Fig 2c**), as they were trained on 2D representations of the chemical molecules (i.e. ECFP4^16^), highlighting the need to develop new descriptors able to distinguish stereoisomer-specific bioactivities.

**Figure 2:**
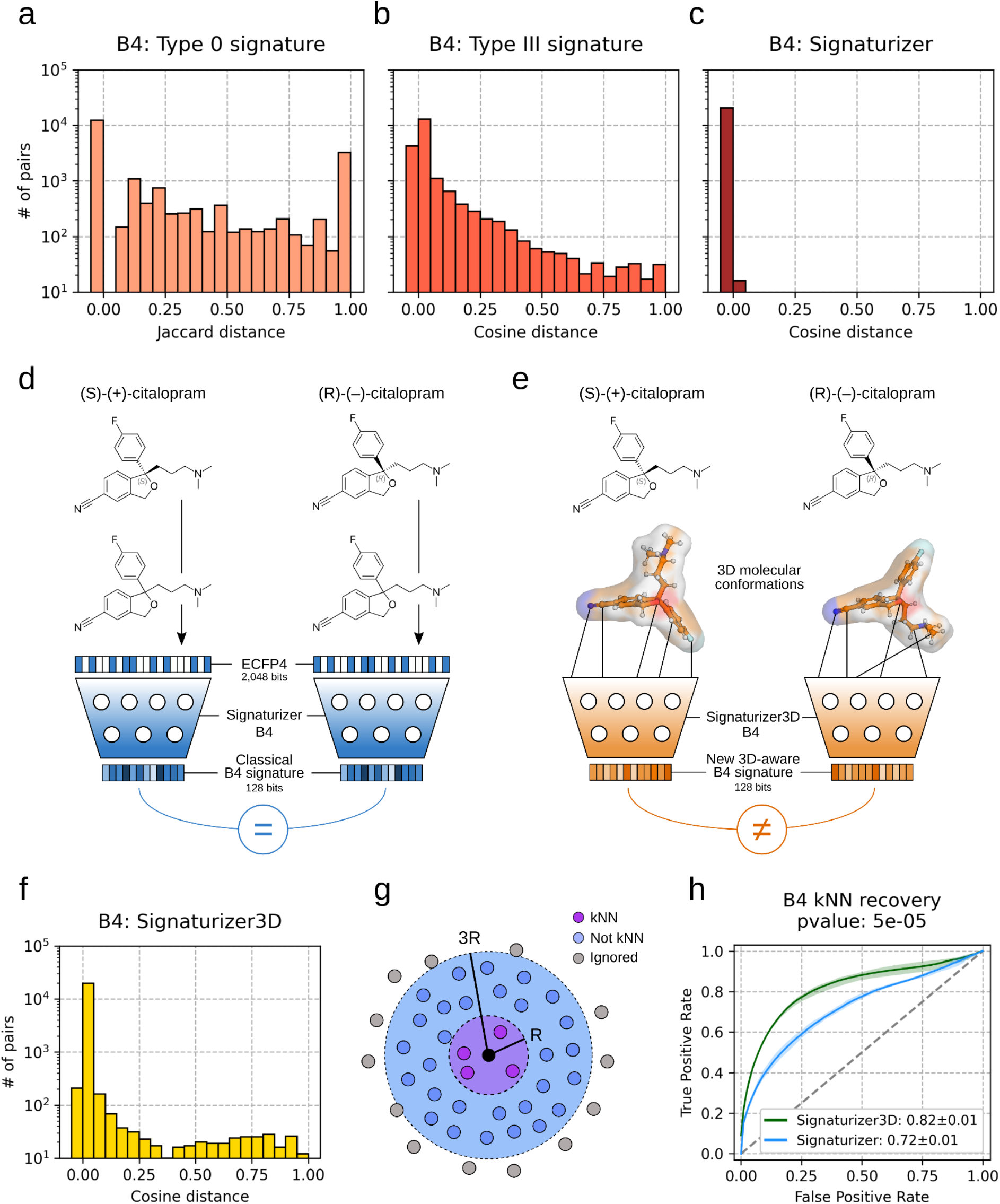
*Stereochemically-aware bioactivity descriptors*. **a)** Distribution of target binding profile Jaccard distances (CC B4 type 0 signatures) between stereoisomer pairs (20,386 pairs). **b)** Distribution of CC B4 type III signature cosine distances between stereoisomer pairs. **c)** Distribution of Signaturizer cosine distances between stereoisomer pairs. **d)** Graphical scheme of the *signaturization* process of distinct stereoisomers ((S)-(+)-citalopram and (R)-(–)-citalopram) with the Signaturizer. Molecules are first represented by 2D-based fingerprints (ECFP4, 3D information is lost) and then input to a neural network. Since ECFP4 for both stereoisomers are identical, output signatures are also identical. **e)** Graphical scheme of the *signaturization* process of distinct stereoisomers ((S)-(+)-citalopram and (R)-(–)-citalopram) with the novel Signaturizer3D. 3D conformations are first generated for both molecules and the corresponding molecular representations are input to the Signaturizer3D fine-tuned neural network. Since molecular representations for both stereoisomers are different, output signatures are also different. **f)** Distribution of Signaturizer3D cosine distances between stereoisomer pairs. **g)** Strategy to assess the recovery of k nearest neighbors (kNN): given a seed compound (black), NNs are defined as those k compounds being at a distance <R (purple). On the other hand, non-NN are defined as those compounds being at a distance >R but <3R (blue). All compounds at a distance >3R (gray) are ignored. **h)** Recapitulation of B4 signature type III kNNs at a p-value ≤ 5x10^-5^ (x3 80/20 splits) using the original Signaturizer and the Signaturizer3D. Positive (NN) and negative (non-NN) pairs were subsampled (1k compounds) from each space as defined in **Fig 2f**.

### Design and evaluation of stereochemically-aware Signaturizers

To overcome the limitations of the original Signaturizers, we trained new deep-networks using 3D-aware molecular representations (i.e. Signaturizers3D, **Fig 2e**). We first generated 3D conformations for all CC molecules, coupled them with their type III signatures, and used them to fine-tune the pre-trained Uni-Mol model^17^ (see *Supplementary Information*). We then evaluated the capability of Signaturizers3D to distinguish stereoisomers by generating B4 signatures for the 32,705 compounds identified in the CC B4 space and calculating distances between stereoisomer pairs. We found that, opposed to the original Signaturizers, virtually all pairs of stereoisomers (99.9%) had non-identical 3D signatures (**Fig 2f**), highlighting the ability of our new models to capture slight differences in the stereochemistry of the compounds. Finally, we followed a strict approach to assess the ability of Signaturizers3D to recapitulate k nearest neighbor (kNN) compounds at type III signature level; this is to evaluate their capacity to retain the structure of the original data similarity. In brief, in a standard kNN recovery task, negative pairs are chosen randomly and can differ significantly from positive pairs. Under this scenario, both Signaturizers and Signaturizers3D could almost perfectly distinguish close from distant molecules at type III signature level (**Fig S2**). To make the assessment more stringent and realistic, we selected the negatives within a close distance of the molecule under evaluation, making the discernment between positive and negative pairs a more difficult task (**Fig 2g**, see Supplementary Information). In this case, we observed that, indeed, Signaturizers3D were able to better recapitulate type III signatures than the original ECFP4-based Signaturizers (**Fig 2h**).

Overall, for the first time, we have systematically assessed the relationship between stereoisomerism and bioactivity on a large scale, focusing on compound-target binding events.

Subsequently, we used our findings to train the second generation of Signaturizers, which are now stereochemically-aware, thereby providing an even more faithful and accurate description of complex small molecule bioactivity properties.

### The Signaturizer3D package

An open source Python package to generate 3D-aware CC bioactivity signatures is available at https://gitlabsbnb.irbbarcelona.org/packages/signaturizer3d. The package includes model weights for each of the 25 CC spaces and can be used to characterize molecules using SMILES or coordinates from existing conformers as input. The models are implemented in Pytorch and support inference on a GPU or CPU. The average time to generate CC signatures from SMILES is 16.3s per 1,000 molecules on an NVIDIA GeForce RTX 3090.

## Acknowledgements

We thank M. Bertoni (Nuage Therapeutics) and M. Duran-Frigola (Ersilia) for thoroughly testing the Signaturizers3D package and for critical reading of the m/s. P.A. acknowledges the support of the Generalitat de Catalunya (2021 SGR 00876), the Spanish Ministerio de Ciencia, Innovación y Universidades (PID2020-119535RB-I00), the Instituto de Salud Carlos III (IMPaCT-Data), and the European Commission (CLARITY: 101137201). A.C-C. is a recipient of an FI fellowship (2020 FI_B 00094). We also acknowledge institutional funding from the Spanish Ministry of Science and Innovation through the Centres of Excellence Severo Ochoa Award, and from the CERCA Programme / Generalitat de Catalunya.

## Author contributions

A.C-C., A.L. and P.A. designed the study. A.C-C., A.L. and M.S. implemented the computational strategy. A.C-C. and P.A. wrote the manuscript. All authors analyzed the results, read and approved the manuscript.

## Conflict of interest

The authors declare no conflict of interest.

## Supplementary Information

### Identification of stereoisomers in the CC universe

We first downloaded small molecule bioactivity data from the official CC website (https://chemicalchecker.com/downloads/signature0, 25 spaces). We standardized a total number of 1,009,293 compounds (InChIs) using the *standardiser* package (https://github.com/flatkinson/standardiser/tree/master, removal of solvent and salt molecules, charge neutralization and application of tautomeric rules), leading to 916,931 standardized and neutral small molecules having 3 or more carbon atoms. Subsequently, we identified groups of stereoisomers based on their InChIKeys^18^, generated using RDKit (https://www.rdkit.org). For compounds to be categorized in the same group, they needed to have identical first blocks (indicating identical connectivities between atoms) and non-identical second blocks (indicating differences in stereochemistry, isotopic atoms, the InChI version, etc). In addition, we excluded small molecules with unspecified chiral centers or unassigned double bond configurations. Finally, we also removed compound pairs having different InChIs after stereochemical information was deleted using RDKit, such as pairs with distinct isotopes. We only conserved groups having ≥2 compounds. Overall, we identified 23,830 groups of stereoisomers, including 57,989 compounds. A graphical scheme of the pipeline is shown in **Fig 1a**.

### Training and evaluation of Signaturizers3D

For all molecules in the CC, we generated and optimized a single 3D conformation per compound using the ETKDG method^19^ and the Merck Molecular Force Field (MMFF94) from RDKit, respectively, with hydrogens included. We successfully generated conformations for ∼99% of the compounds (998,186). The rest, mostly very large molecules, were excluded from the final dataset. We then did 80/20 scaffold-based splits (x3) of the 998,186 considered compounds.

After removing hydrogens, all coordinates and atom-types for each molecule were used to fine-tune the pre-trained Uni-Mol model as a multitarget regression problem, so that we could directly infer pre-calculated CC type III signatures (128 dimensions). Uni-Mol pre-trained model was directly downloaded from https://github.com/dptech-corp/Uni-Mol/releases/download/v0.1/mol_pre_no_h_220816.pt, which provided the initial weights and defined the model’s architecture. To fine-tune the existing Uni-Mol model, we trained for 40 epochs with a batch size of 32 and a learning rate of 0.0001 (warmup ratio of 0.06 without dropout). We used the Adam optimizer with betas (0.9, 0.99) and epsilon 10^-6^, and we utilized a polynomial decay scheduler for the learning rate. Models were trained on the previously generated scaffold-based splits (3x25, 80/20) using a smooth mean absolute error loss for function optimization. Validation was based on the aggregated mean absolute error (*valid_agg_mae*), with early stopping after 20 epochs if no improvement was observed. Training was executed on a GPU with mixed precision (FP16).

### K nearest neighbors recovery

First, we needed to establish a distance cutoff to eventually define positive pairs of compounds (nearest neighbors) at type III signature level. To do so, we sampled 10k molecules in each of the 25 bioactivity spaces and, after calculating the full pairwise distance matrix between these compounds (10^8^ distances within each space), we established the NN cutoff distance as the 0.00005 percentile of the distribution. In this way, a distance cut-off was defined for each CC space.

Positive pairs (NN) for a given compound at type III signature level were defined as those compounds being at a distance lower than the established cutoff for each CC space. For the strict NN recovery, negative pairs (not NN) were defined as those being at a distance greater than the established cutoff but lower than three times the cutoff distance. The ratio of negative to positive pairs was capped at 10:1. We calculated distances between compounds using Signaturizers and Signaturizers3D for all (x3) scaffold-based 80/20 splits, and we assessed their ability to recapitulate NNs at type III signature level by calculating the corresponding ROC Curves with a subsample of 1k molecules per space.

**Figure S1:**
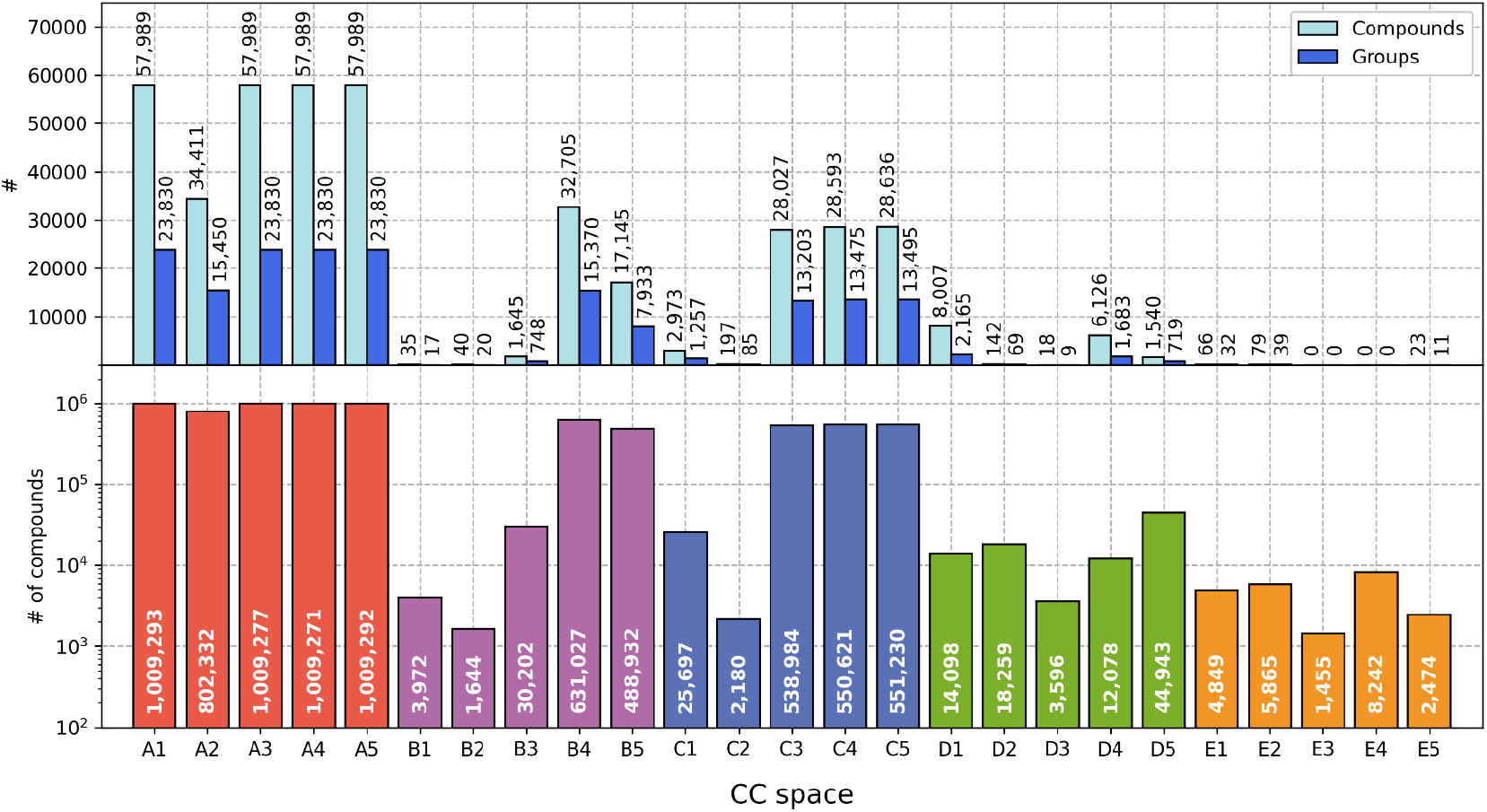
Number of compounds (bottom) in each CC space. Number of groups of stereoisomers and number of stereoisomers (top) in each CC space.

**Figure S2:**
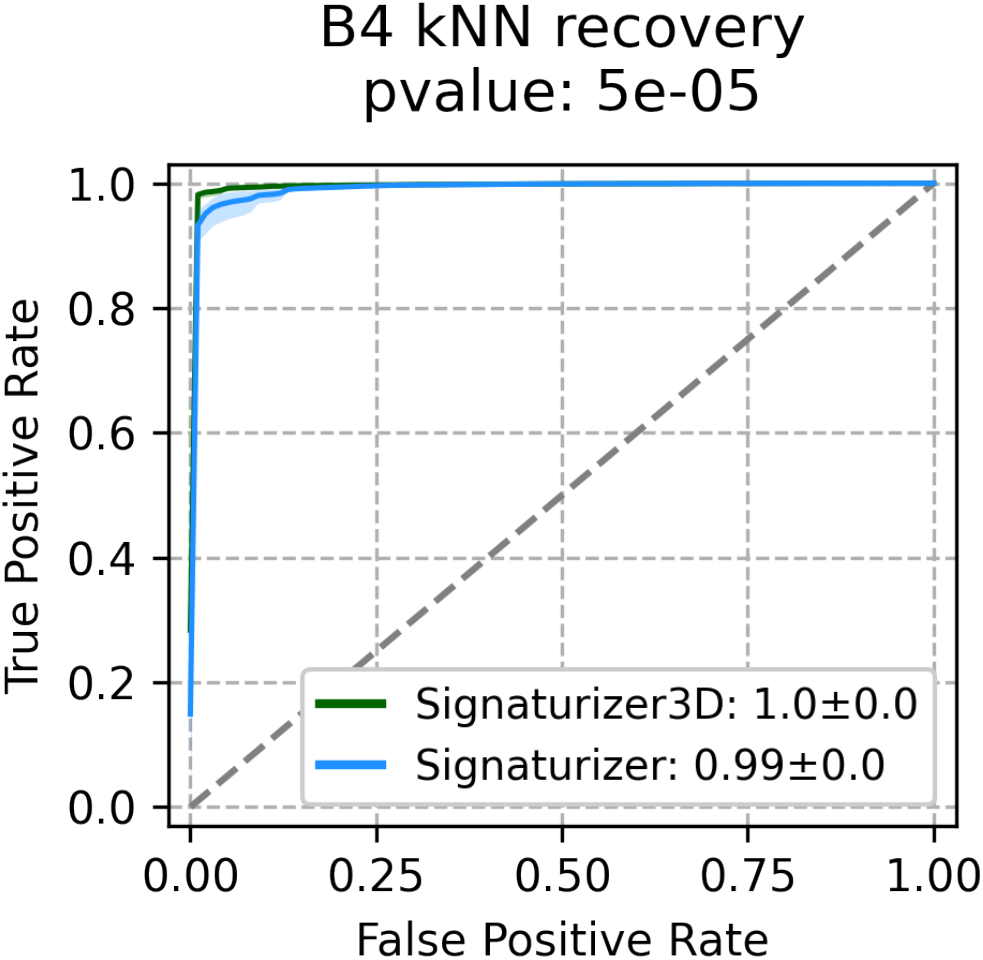
Recapitulation of B4 signature type III kNNs at a p-value ≤ 5x10^-5^ (x3 80/20 splits) using the original Signaturizer and the Signaturizer3D. As opposed to **Fig 2g** and **Fig 2h**, negative pairs were randomly subsampled from all compounds being at a distance above the established threshold (>R).

## Notes

https://gitlabsbnb.irbbarcelona.org/packages/signaturizer3d

### Competing Interest Statement

The authors have declared no competing interest.

## References

1. Bertoni, M. et al. Bioactivity descriptors for uncharacterized chemical compounds. Nat Commun 12, 3932 (2021).

2. Zdrazil, B. et al. The ChEMBL Database in 2023: a drug discovery platform spanning multiple bioactivity data types and time periods. Nucleic Acids Res 52, D1180–D1192 (2024).

3. Kim, S. et al. PubChem 2023 update. Nucleic Acids Res 51, D1373–D1380 (2023).

4. Tetko, I.V., Engkvist, O., Koch, U., Reymond, J.L. & Chen, H. BIGCHEM: Challenges and Opportunities for Big Data Analysis in Chemistry. Mol Inform 35, 615–621 (2016).

5. von Lilienfeld, O.A. & Burke, K. Retrospective on a decade of machine learning for chemical discovery. Nat Commun 11, 4895 (2020).

6. Fernández-Torras, A., Comajuncosa-Creus, A., Duran-Frigola, M. & Aloy, P. Connecting chemistry and biology through molecular descriptors. Current Opinion in Chemical Biology 66, 102090 (2022).

7. Duran-Frigola, M. et al. Extending the small-molecule similarity principle to all levels of biology with the Chemical Checker. Nat Biotechnol 38, 1087–1096 (2020).

8. Scott, K.A. et al. Stereochemical diversity as a source of discovery in chemical biology. Current research in Chemical Biology 2, 100028 (2022).

9. Brooks, W.H., Guida, W.C. & Daniel, K.G. The significance of chirality in drug design and development. Curr Top Med Chem 11, 760–770 (2011).

10. Inaki, M., Liu, J. & Matsuno, K. Cell chirality: its origin and roles in left-right asymmetric development. Philos Trans R Soc Lond B Biol Sci 371 (2016).

11. McConathy, J. & Owens, M.J. Stereochemistry in Drug Action. Prim Care Companion J Clin Psychiatry 5, 70–73 (2003).

12. Smith, S.W. Chiral toxicology: it’s the same thing…only different. Toxicol Sci 110, 4–30 (2009).

13. Sanchez, C., Bogeso, K.P., Ebert, B., Reines, E.H. & Braestrup, C. Escitalopram versus citalopram: the surprising role of the R-enantiomer. Psychopharmacology (Berl) 174, 163–176 (2004).

14. Sanchez, C. The pharmacology of citalopram enantiomers: the antagonism by R-citalopram on the effect of S-citalopram. Basic Clin Pharmacol Toxicol 99, 91–95 (2006).

15. Williams, K.M. Enantiomers in arthritic disorders. Pharmacol Ther 46, 273–295 (1990).

16. Rogers, D. & Hahn, M. Extended-connectivity fingerprints. J Chem Inf Model 50, 742–754 (2010).

17. Zhou, G. et al. Uni-Mol: a universal 3D molecular representation learning framework. ChemRxiv (2022).

18. Heller, S.R., McNaught, A., Pletnev, I., Stein, S. & Tchekhovskoi, D. InChI, the IUPAC International Chemical Identifier. J Cheminform 7, 23 (2015).

19. Riniker, S. & Landrum, G.A. Better Informed Distance Geometry: Using What We Know To Improve Conformation Generation. J Chem Inf Model 55, 2562–2574 (2015).

